# PI3K/AKT Signaling Mediates Stress-Inducible Amyloid Aggregation Through c-myc

**DOI:** 10.1101/2024.09.05.611538

**Authors:** Emma Lacroix, Evgenia A. Momchilova, Sahil Chandhok, Mythili Padavu, Richard Zapf, Timothy E. Audas

## Abstract

In response to environmental stress, eukaryotic cells reversibly form functional amyloid aggregates, called amyloid bodies (A-bodies). While these solid-like biomolecular condensates share many biophysical characteristics with pathological amyloids, A-body are non-toxic, and induce a protective state of cellular dormancy. As a recently identified structure, the modulators of A-body biogenesis remain uncharacterized, with the seeding noncoding RNA being the only known regulatory factor. Here, we use an image-based high-throughput screen to identify candidate pathways regulating A-body biogenesis. Our data demonstrates that the PI3K signaling axis meditates A-body formation during heat shock, by activating AKT and repressing GSK3-mediated degradation of c-myc. This enhances c-myc binding to regulatory elements of the seeding noncoding RNA, upregulating the transcripts that nucleate A-body formation. Identifying a link between PI3K signaling, c-myc, and physiological amyloid aggregates, extends the range of activity for these well-established regulators, while providing insight into cellular components whose dysregulation could underly amyloidogenic disorders.

## Introduction

In response to extracellular signals, cells alter their metabolic behaviour^1^. This modulation is usually achieved through the activation of signaling cascades, where the sensing of fluctuating conditions is transmitted to specific downstream effectors^2^. The binding of ligands to membrane-associated receptors often initiates this process, but it is the activation of a complex web of intracellular signaling molecules that leads to the alteration of critical cellular processes, such as proliferation, migration, and metabolism^3–5^. Harsh environmental stimuli (e.g., heat, acidity, and DNA damage) can also activate signaling cascades, a process that is collectively referred to as the cellular stress response. However, different stressors will cause different types of damage, necessitating a carefully tailored response to the unique perturbations. Pro-survival pathways will attempt to repair the damage, but programmed cell death may ultimately be needed to maintain viability of the organism^6–10^.

In recent years, biomolecular condensates have emerged as stress responsive hubs within eukaryotic cells. Membrane-less organelles are ideally positioned for this task, as these structures can rapidly assemble and disassemble in response to changing environmental conditions^11–13^. Stress-induced signal transduction events have been shown to control this process, as they can alter the abundance of scaffolding molecules that mediate condensate stability^14–16^. When present, these structures modulate the biological activity of their resident biomolecules. Critical regulators can be inhibited by sequestration away from their downstream molecular networks or reaction kinetics can be enhanced by micro-concentrating cellular factors together within a specific subcellular region. For example, cytoplasmic stress granules sequester mRNA away from ribosomes to downregulate translation under various stress conditions^17,18^, while heat-induced nuclear stress bodies micro-concentrate transcription and splicing machinery to enhance the expression and maturation of specific genes^14,19^. As conditions change, the lack of a membrane allows these condensates to rapidly exchange material with the cellular milieu, further enhancing the adaptability and versatility of these stress-responsive hubs^12,20^.

Nucleoli are particularly intriguing biomolecular condensates, as they have a role in both basal and stress-responsive pathways^21^. Under normal growth conditions the nucleolus transcribes and processes RNA for ribosome assembly^22–24^; however, temperature- and pH-sensitive signaling drastically alters the structure and function of this condensate. Here, stress treatment represses ribosome biogenesis, while concomitantly enhancing expression of noncoding RNA (ncRNA) derived from the ribosomal intergenic spacer (rIGS)^25–27^. Each stressor stimulates the transcription of a different locus of the rDNA cassette, with acidosis mediating expression of a sequence ∼28kb downstream of the ribosomal rRNA transcriptional start site (rIGS_28_RNA) and heat shock exposure upregulating the 16kb and 22kb regions (rIGS_16_RNA and rIGS_22_RNA)^27^. Despite these differing loci, each rIGSRNA transcript contains repetitive and low complexity sequences, which enables the ncRNAs to mediate the recruitment of a wide array of cellular proteins through complex coacervation^25^. Following a short lag phase, the recruited proteins coalesce within this subnuclear region, replacing the classic tripartite nucleolar structure with large, dense, and fibrillar aggregates^25,28–34^. A biophysical assessment of these stress-induced aggregates revealed that the resident proteins assemble into highly-ordered amyloid fibrils^28^, which share the hallmark characteristics of amyloid plaques commonly observed in neurodegenerative disorders (e.g., Alzheimer’s, Parkinson’s, and prion-based diseases)^35–37^. To acknowledge the complete dissolution of the nucleolar architecture, the foci were named Amyloid bodies (A-bodies), and they joined a growing list of functional amyloids that have been uncovered in the literature^38–40^. Biologically, the formation of A-bodies induces a state of cellular dormancy, which promotes cell viability during exposure to harsh environmental stimuli^28^, demonstrating that A-body biogenesis is an important component of the cellular stress response.

While the role of the regulatory ncRNA that mediate A-body formation has been characterized^25,27,28^, no other components of the signaling axis for these stress-induced functional amyloids have been identified. Thus, we performed a high-throughput image-based analysis to identify sensors, signals, and effectors that promote A-body formation. By screening a collection of 4344 chemical compounds that inhibit or activate a variety of known cellular proteins, we identified 147 small molecules that repress A-body biogenesis. As many of these compounds targeted PI3K signaling pathway members, we demonstrated that this signaling network is strongly activated by high temperatures and needed for the optimal formation of A-bodies. By mapping downstream pathway components, we found c-myc to be a critical effector of PI3K signaling in heat shock. This proto-oncogenic transcription factor associates with the rIGS_16_RNA locus, and promotes expression of the regulator ncRNA in a PI3K-dependent manner. Together, these data highlight a role for the PI3K pathway and c-myc in the formation of physiological amyloid aggregates.

## Results

### The PI3K Signaling Cascade Promotes A-body Formation

Harsh stimuli can activate an array of signaling networks that drive stress-specific cellular responses. To date, no signaling molecules have been identified that mediate A-body biogenesis, leaving this amyloid aggregation pathway poorly characterized. To begin identifying these upstream regulators, we developed a high-throughput image-based screening approach, which monitors the localization of a GFP-tagged A-body targeting sequence (ATS-GFP)^28^ that can be reversibly sequestered within these condensates in response to multiple stressors (**Figure 1A**, **Figure S1A**). Cells stably-expressing this reporter construct were pre-treated for 24-hours with a diverse collection of 4344 bioactive compounds, which target molecules involved in a variety of cellular pathways including; membrane transport, inflammatory response, cell cycle progression, metabolism, and apoptosis. Following a heat shock treatment, the cells were fixed, imaged, and scored (**Figure S1B**), to identify 147 bioactive compounds that putatively reduce A-body aggregation by at least 3 standard deviations of the mean (**Figure 1B**, **Figure 1C - purple bars, Figure 1D**). To begin validating these results, we individually purchased a subset of the candidate compounds, and quantitatively assessed their effect on A-body recruitment using a previously established methodology (**Figure 1E**)^29,31,32^. Our low-throughput data corroborated the screening results, demonstrating a significant reduction in heat-induced A-body recruitment from a number of the putative hits (**Figure 1E**).

**Figure 1.**
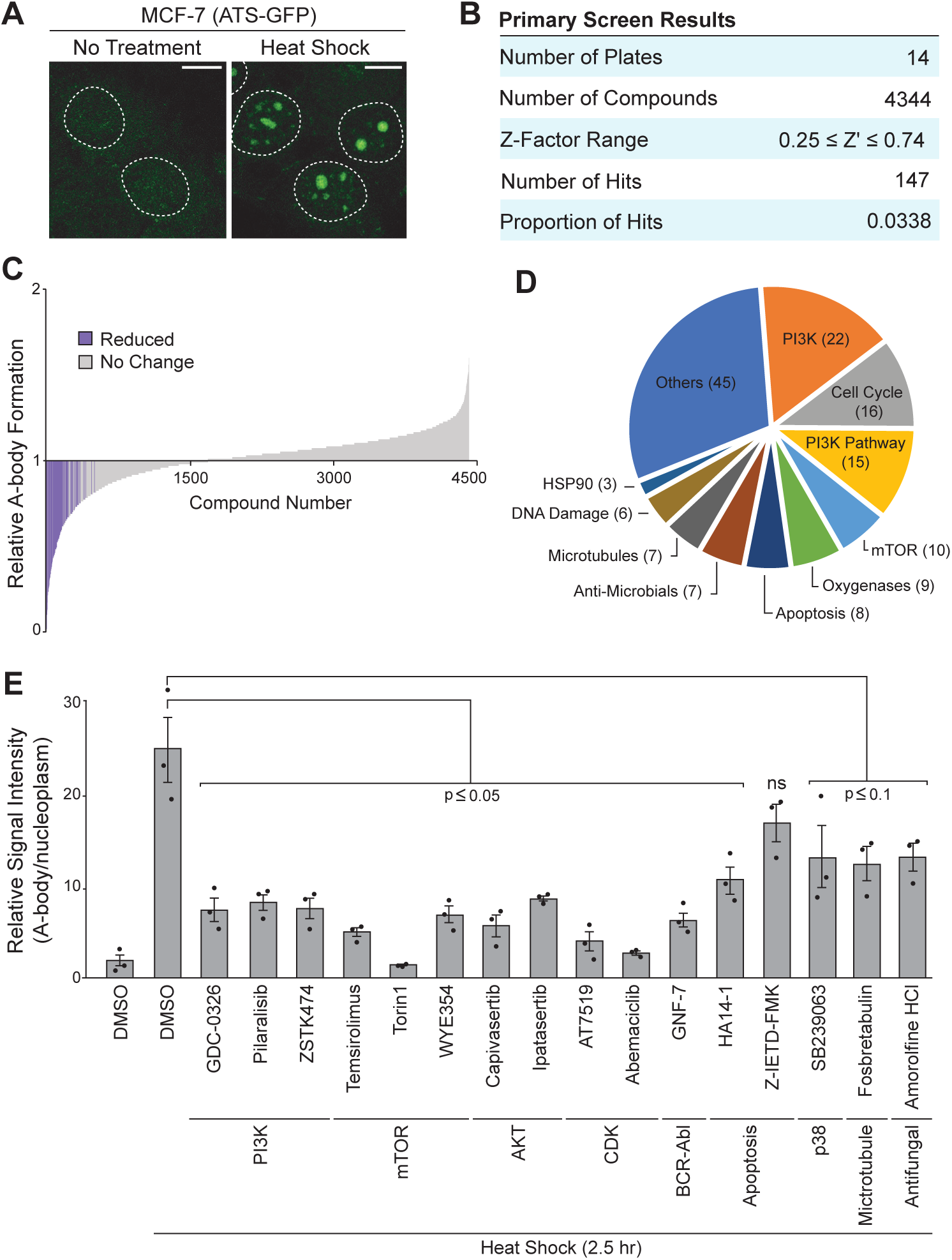
High-Throughput Image-Based Screen Reveals Cellular Factors that Regulate A-body Formation. **(A)** MCF-7 cells stably expressing the ATS-GFP reporter construct were left untreated or exposed to a 2-hour heat shock treatment (43°C). Dashed white lines represent nuclei. Scale bars: 10µm. **(B)** Summary of the High-Throughput screening results. **(C)** Relative A-body formation was quantified for all compounds in our screen that met our selection criteria. Putative hits (purple) are greater than 3 standard deviations below the negative control. **(D)** Pathway analysis of screening hits. Pathways with 3 or more compounds that reduced A-body formation are listed. **(E)** The MCF-7 (ATS-GFP) cell line was treated with the indicated compounds (20µM) for 24-hours, prior to heat shock exposure (2.5 hours). Relative A-body/nucleoplasmic fluorescence intensity was calculated for each sample. Individual data points are included on the graph (black dots), and values represent means ± SEM (n=3).

Nearly half of the candidate compounds target either proteins in the PI3K/mTOR/AKT signaling axis or members of the cyclin-dependent kinase (CDK) family (**Figure 1D**). Interestingly, CDK inhibition was recently shown to disrupt basal nucleolar architecture^41^, which is the progenitor structure that is re-modeled into A-bodies during periods of cellular stress. As the effect of the CDK inhibitors could be tied to impaired nucleolar morphology (**Figure S2A**), rather than inactivation of the A-body assembly machinery, we focused our study on the role of PI3K/mTOR/AKT signaling in amyloid aggregate formation. Class 1 PI3Ks are lipid kinases that catalyze the formation of phosphatidylinositol-3,4,5-trisphosphate within the plasma membrane^42,43^. Once modified, these lipids act as a docking site for AKT^42,44,45^, which can then be activated by the kinases PDK1 and mTORC2, who phosphorylate critical threonine 308 (T308) and serine 473 (S473) residues, respectively^46^. This signaling cascade then modulates a vast network of downstream effectors, which are associated with a broad array of cellular pathways including proliferation, survival, and protein synthesis^47^. For the PI3K/mTOR/AKT pathway to play a role in this amyloid aggregation process, it would need to be active at the same temperatures where A-body biogenesis occurs. Using an antibody to detect active AKT (phosphorylated S473), we performed a thermo-course experiment and found elevated AKT phosphorylation at ∼42-43°C, which tightly paralleled the A-body formation kinetics seen with the reporter ATS-GFP construct (**Figure 2A-B**). Next, we shortened the compound pre-treatment time (1-hour) to demonstrate that the observed effects were associated with target molecule inhibition, rather than the result of downstream signaling events or cellular adaption that can occur upon prolonged (24-hour) drug exposure. This shorter treatment course was sufficient to disrupt AKT1 activation (**Figure S2B**), and we found that PI3K, mTOR, and AKT1 inhibitors continued to impair ATS-GFP aggregation under these conditions (**Figure 2C**). To confirm that these signaling inhibitors were broadly disrupting A-body formation, and not just targeting of the artificial reporter construct, we monitored the recruitment of endogenous A-body constituents CDC73 and MDM2 and the binding of Congo red (**Figure 2D**). The significant reduction in Congo red signal, was particularly notable as this amyloidophilic dye detects any proteins that have adopted the amyloid conformation, demonstrating that A-body formation was broadly inhibited by each of these compounds (**Figure 2D-E**). It is important to note that most of the mTOR hits found in our screen were dual mTORC1 and mTORC2 inhibitors (**Figure 1**); thus, we tested the mTORC1-specific inhibitor rapamycin to distinguish between the activity of the two complexes. Though the mTORC1/2 dual inhibitor Torin1 reduced A-body formation significantly (**Figure 2C-E**), rapamycin did not (**Figure 2E**), suggesting that mTORC2 is likely responsible for this effect. To verify that these results were not due to off-target effects of the small molecules, we depleted mTOR and AKT1 using siRNAs (**Figure S2C, Figure S2D**). These depletions resulted in a significant reduction in the number of cells containing A-bodies (**Figure S2E, Figure S2F**), supporting a role for the PI3K/mTORC2/AKT signaling axis in A-body biogenesis.

**Figure 2.**
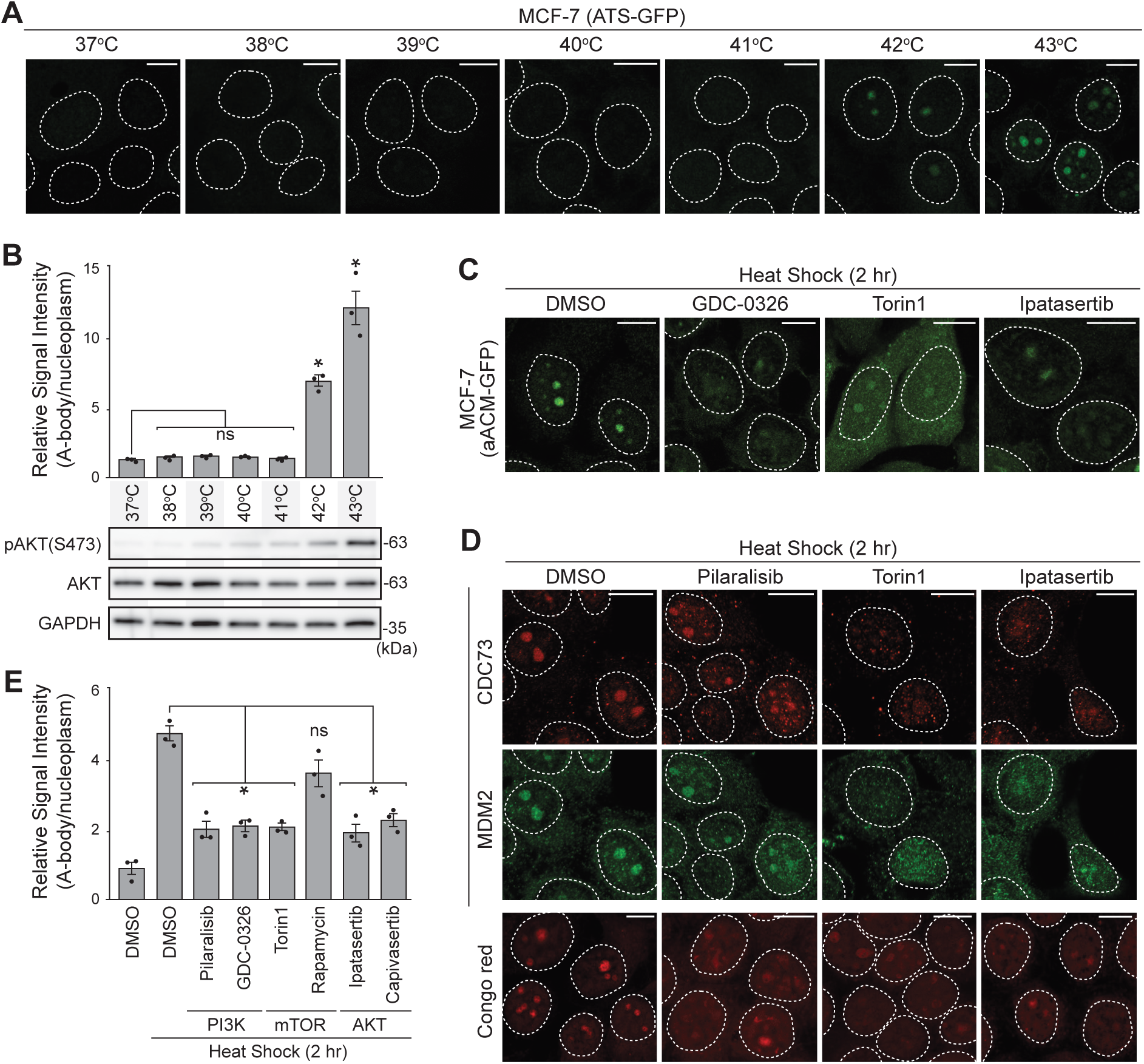
The PI3K Signaling Cascade Promotes A-body Formation. **(A)** MCF-7 (ATS-GFP) reporter cells were exposed to the indicated temperatures for 2 hours and representative images were captured. **(B)** Quantification of ATS-GFP signal in the A-bodies, relative to the nucleoplasm, was calculated for the samples in (A) (top). Western blotting to detect AKT, phosphorylated AKT (S473), and GAPDH was performed on MCF-7 cells exposed to the indicated temperatures for 30 minutes (bottom). **(C)** MCF-7 (ATS-GFP) cells were pre-treated with the indicated inhibitors (20µM) for 1 hour, then heat shocked at 43°C for 2 hours. **(D)** MCF-7 cells pre-treated for 1 hour with 20µM of the indicated compounds were heat shocked for 2 hours prior to either immunostaining for endogenous CDC73 and MDM2 (top) or amyloidophilic dye staining with Congo red (bottom). **(E)** Heat shock exposed (2 hours) MCF-7 cells were pre-treated (1 hour) with DMSO or the indicated PI3K, mTOR, and AKT inhibitors (20µM). Following Congo red staining, the signal intensity of the A-bodies was quantified relative to the background nuclear fluorescence. Dashed white circles represent nuclei, and scale bars are 10µm. Individual data points are included on the graph (black dots), and values represent means ± SEM (n=3). *p≤0.05 (two-tailed unpaired student’s t-test).

### PI3K/mTORC2/AKT Signaling Regulates rIGS_16_RNA Expression by Inhibiting GSK3α/β

Upregulation of ncRNA expressed from the ribosomal intergenic spacer is the earliest know event in A-body biogenesis, though cellular regulators of these noncoding transcripts have not been established^25,27,28^. By comparing the temporal induction of the rIGS_16_RNA and the heat-induced activation of the PI3K/mTORC2/AKT signaling pathway, we found that AKT phosphorylation occurred prior to the expression of this ncRNA transcript (**Figure 3A**). Therefore, we hypothesized that PI3K signaling was promoting A-body formation through the expression of rIGS_16_RNA. Using qPCR, we found that inhibitors of PI3K, mTORC1/2, and AKT all significantly reduced heat-mediated induction of rIGS_16_RNA (**Figure 3B**), without affecting GAPDH mRNA levels (**Figure S3A**). The same outcome was observed with siRNA-mediated depletion of AKT1 (**Figure 3C**), together demonstrating a role for PI3K signaling in the induction of rIGS_16_RNA expression. To help identify downstream targets of AKT1 that could promote the induction of rIGS_16_RNA, we returned to the high-throughput screening results. Performing an alternate analysis of the data, for compounds that enhance A-body formation, we found 66 hits that increased A-body biogenesis by at least 3 standard deviations (**Figure S3B**). This list of A-body enhancers included inhibitors of GSK3α/β, as well as a number of other cellular pathways (**Figure S3C**). GSK3α/β are constitutively active kinases, whose activity is directly inhibited by AKT phosphorylation^48,49^. Thus, impairment of GSK3α/β should theoretically give the opposite phenotype to AKT inhibitors, and result in the enhancement of A-body biogenesis. Two GSK3α/β inhibitors (LY2090314 and CHIR-98014) were purchased, and their effects on A-body formation was assessed. To enhance our ability to detect this putative increase in A-body biogenesis, we reduced the heat shock treatment time to 1 hour, which is well below the time when Congo red signal intensity reaches saturation. Under these conditions, we found that GSK3α/β inhibition significantly increased A-body biogenesis (**Figure 3D-E**), and enhanced CDC73 and MDM2 recruitment within these amyloid-based structures (**Figure 3F**). Consistent with the temporal induction of AKT1 (**Figure 3A**), inhibitory phosphorylation events were observed on GSK3α and GSK3β within 15 minutes of heat shock exposure (**Figure 3G**), though this could be impaired by treatment of cells with PI3K, mTORC1/2, and AKT inhibitors (**Figure 3H**). Examining the heat-induced upregulation of rIGS_16_RNA expression, we found that the levels of this ncRNA were significantly enhanced by GSK3α/β inhibitor treatment (**Figure 3I**), without affecting GAPDH mRNA expression (**Figure S3D**). Conversely, over-expression of exogenous GSK3α/β impaired the reduction of rIGS_16_RNA that was mediated by AKT inhibition (**Figure S3E**). This PI3K/mTORC2/AKT/GSK3α/β- mediated regulation of A-body formation/rIGS_16_RNA expression was also observable in U-87 MG cells, demonstrating that this effect is not cell-line specific (**Figure S3F, Figure S3G**). Together, these data suggest that PI3K/mTORC2/AKT-mediated GSK3α/β inactivation promotes rIGS_16_RNA expression to regulate the formation of A-bodies during heat shock exposure.

**Figure 3.**
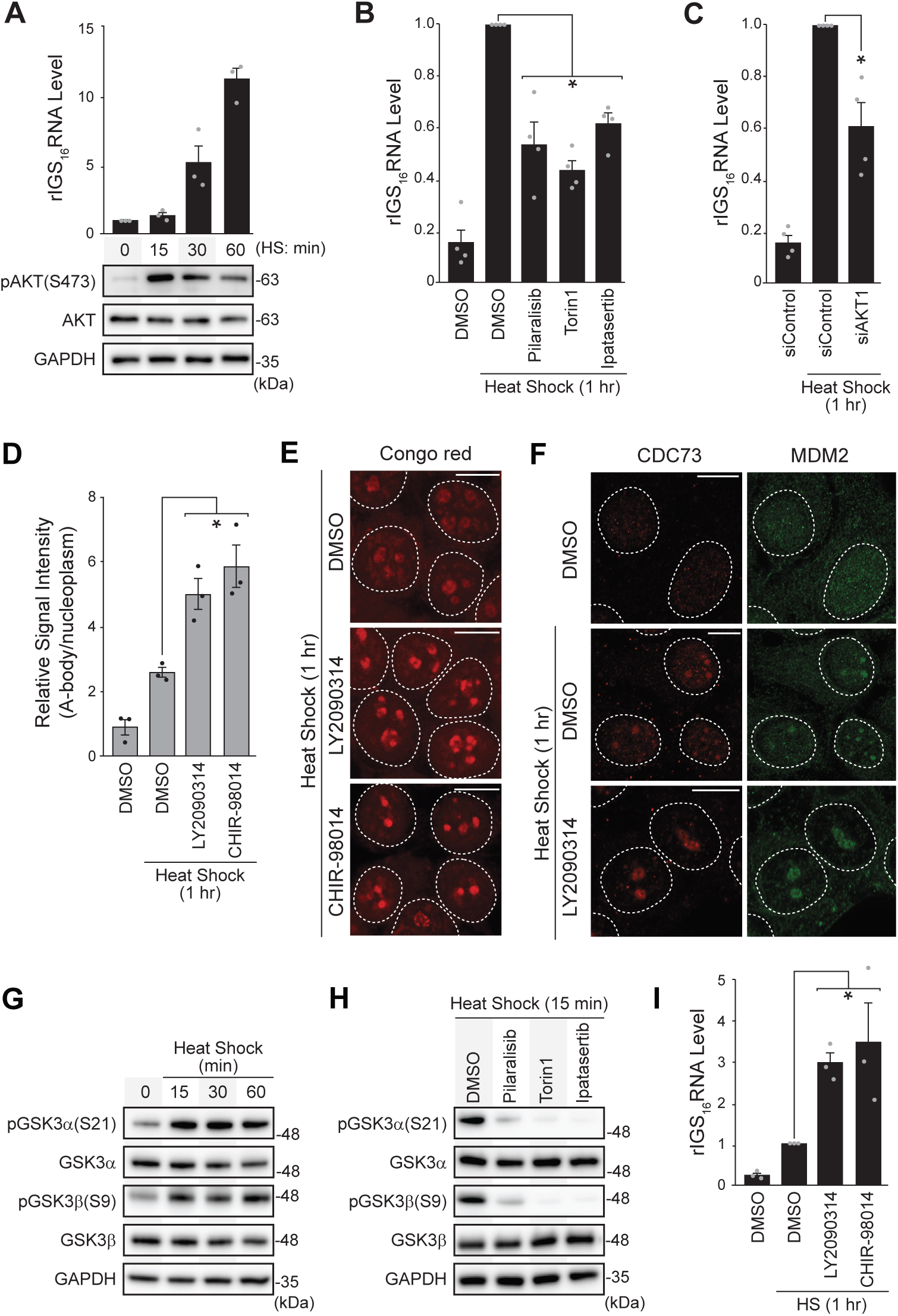
PI3K/mTORC2/AKT Signaling Regulates rIGS_16_RNA Expression by Inhibiting GSK3α/β. **(A)** MCF-7 cells were heat shock treated at 43°C for the indicated times. (Top) Quantitative PCR was used to calculate the rIGS_16_RNA levels. (Bottom) Western blotting detected AKT activation, by assessing total AKT, phosphorylated AKT (S473), and GAPDH protein levels under the same conditions. **(B)** The effects of chemical inhibition (20µM, 1-hour pre-treatment) of PI3K (Pilaralisib), mTOR (Torin1), and AKT (Ipatasertib) on rIGS_16_RNA expression by qPCR in cells heat shock treated for 1 hour. **(C)** Quantification of rIGS_16_RNA levels in siAKT1 transfected cells (30pmol, 48-hours), heat shock treated for 1 hour at 43°C. **(D, E)** The GSK3 inhibitors LY2090314 and CHIR-98014 were added (20µM) 1 hour prior to heat shock treatment. (D) Relative A-body/nucleoplasmic signal was quantified for Congo red stained MCF-7 cells. (E) Representative images are included. **(F)** Endogenous CDC73 and MDM2 were detected by immunostaining of untreated or heat shock treated (1 hour) MCF-7 cells pre-treated with DMSO or LY2090314 (20µM). **(G, H)** GSK3α/β phosphorylation levels were assessed by western blotting samples with antibodies targeting GSK3α, phosphorylated GSK3α (S21), GSK3β, phosphorylated GSK3β (S9), and GAPDH. (G) MCF-7 cells were exposed to 43°C conditions for the indicated times, or (H) heat shock treated (15 min) cells were pre-treated (1 hour) with the indicated PI3K pathway inhibitors (20µM). **(I)** rIGS_16_RNA levels were quantified by qPCR from untreated or heat shock treated (1 hour) MCF-7 cells pre-treated with DMSO or the GSK3 inhibitors LY2090314 and CHIR-98014 (20µM). RNA expression qPCR data is normalized to β-actin levels. Individual data points are included on the graphs (black or grey dots) and values on graphs represent means ± SEM (n≥3). *P≤0.05 (two-tailed unpaired student’s t-test). Dashed white circles in microscopy images represent nuclei. Scale bars: 10µm.

### c-myc Accumulates in the Nucleolus during Heat Shock to Induce rIGS_16_RNA Expression

GSK3α and GSK3β are serine/threonine kinases that were initially identified through their role in glycogen synthesis^50^, though subsequent work has shown that transcription factors are their largest group of substrates^49^. As rIGS_16_RNA upregulation is likely a transcriptional event, we explored the high-throughput screening results for transcription factors that are known to be downstream targets of GSK3α/β. Interestingly, our data contained a single hit compound that targeted c-myc, a transcription factor that binds to E-box elements within DNA to enhance the expression of over 15% of the genome^51^. Degradation of this proto-oncogene is regulated by GSK3α/β-mediated phosphorylation^49,52^, suggesting that inactivating these kinases would increase the transcriptional activity of c-myc protein. To explore the role of c-myc on A-body formation, we first needed to test if this transcription factor could be found at the site of rIGS_16_RNA expression, the nucleolus^25–27^. During heat shock exposure c-myc rapidly accumulates in the nucleolus (**Figure 4A**). This notably occurs prior to the heat-induced expression of rIGS_16_RNA (**Figure 3A**), highlighting that the nucleolar localization of this transcription factor was not mediated by the rIGSRNA itself, as these ncRNA species have been shown to recruit A-body resident proteins^25,27^. Depletion of c-myc using siRNA (**Figure S4A**), significantly reduced both A-body formation (**Figure 4B-C**), and the levels of rIGS_16_RNA expression under heat shock conditions (**Figure 4D**), without affecting GAPDH (**Figure S4B**), suggesting that c-myc activity is involved in this amyloid aggregation pathway. To explore whether this c-myc-mediated effect was linked to the PI3K signaling axis, we assessed c-myc localization in cells treated with PI3K pathway inhibitors and found that c-myc levels appeared to be reduced throughout the nucleus/nucleolus (**Figure 4E**). Next, we analyzed c-myc transcript and protein levels during heat shock exposure. Despite the mRNA remaining constant (**Figure S4C**), protein abundance steadily increased over the course of a 1-hour heat shock treatment (**Figure 4F**). As predicted, the PI3K inhibitor (Pilaralisib) reduced c-myc protein levels, while inactivation of GSK3α/β (LY290314) resulted in further accumulation of the protein (**Figure 4G**). These observations were paralleled in U-87 MG cells (**Figure S4D-E**), indicating that this form of c-myc regulation was not cell line specific. As it has been previously shown that c-myc binds E-box elements in the rDNA cassette to promote ribosomal RNA expression^53,54^, we explored whether this transcription factor could also associate with E-box elements near the rIGS_16_ genomic locus to mediate heat shock-induced expression of the ncRNA (**Figure 4H**). Chromatin immunoprecipitation (ChIP) data supports this notion, as unlike other c-myc-driven promoters (UCP2 and MXD3) or an unrelated region of the rDNA cassette (rIGS_28_), endogenous c-myc binding to the rIGS_16_RNA-encoding region was significantly enhanced at high temperatures (**Figure 4I, Figure S4F**). Additionally, this heat-induced association was impeded by PI3K inhibition (**Figure 4I, Figure S4F**), highlighting a role for the PI3K signaling axis in this c-myc-driven transcriptional event. Taken together, our data demonstrates that PI3K/mTORC2/AKT-mediated inactivation of GSK3α/β is responsible for stabilizing c-myc protein in heat shock-treated cells. This leads to enhanced c-myc binding of the rIGS_16_RNA locus, and upregulation of ncRNA that drive physiological amyloid aggregation.

**Figure 4.**
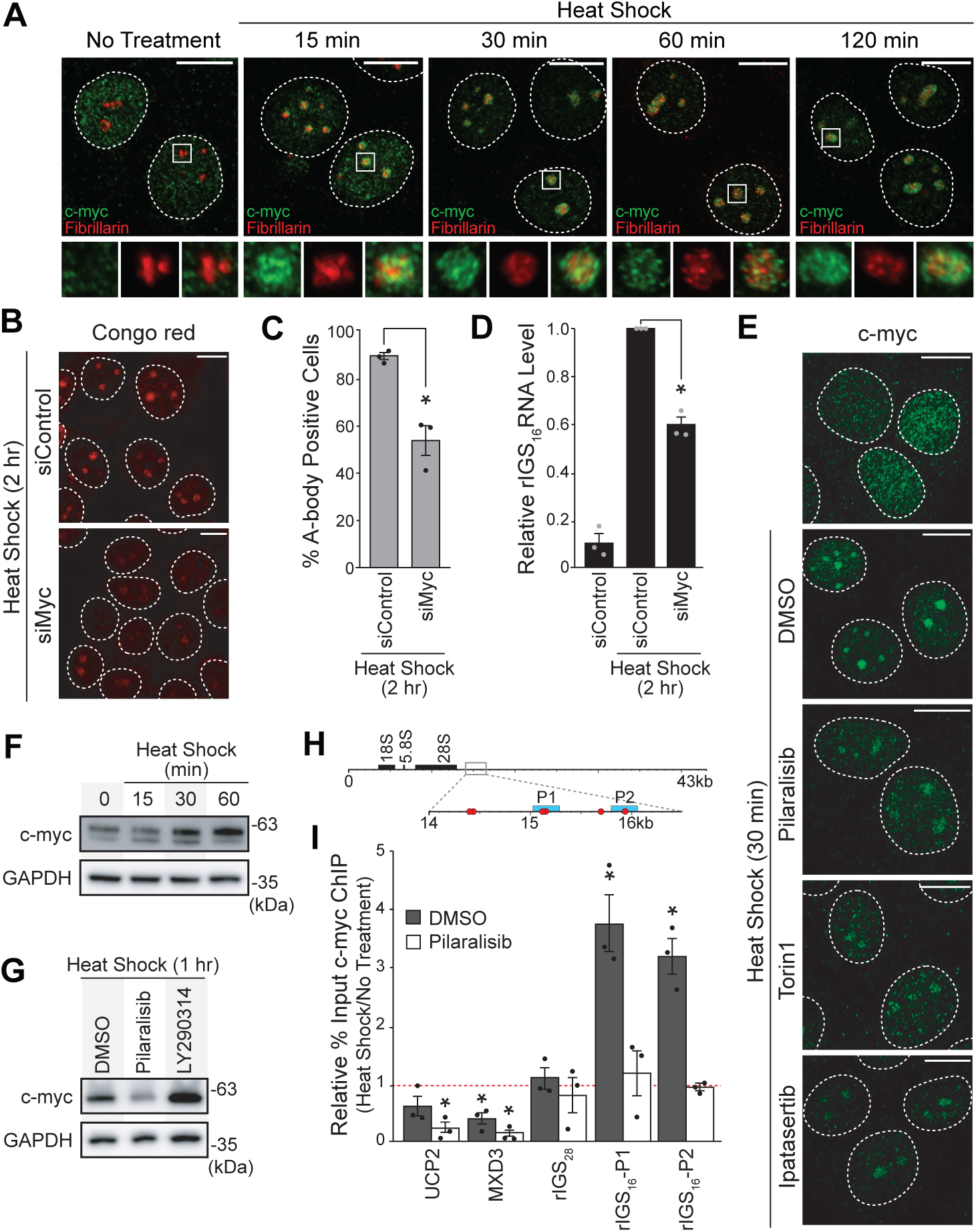
c-myc Accumulates in the Nucleolus during Heat Shock to Induce rIGS_16_RNA Expression. **(A)** MCF-7 cells were heat shock treated for the indicated times, prior to immunodetection with c-myc (green) and fibrillarin (red) antibodies. White square insert is magnified below, with merge panel on the right. **(B-C)** An siRNA-mediated knockdown of c-myc (30pmol, 48-hours) was performed in MCF-7 cells. (B) Representative images of heat shock-treated (2 hours) cells stained with Congo red were captured, and (C) the percentage of A-body-positive cells was calculated. A minimum of 500 cells across 3 independent biological replicates were scored as either A-body-positive or -negative. **(D)** Knockdown of c-myc was performed as in (B) and rIGS_16_RNA expression was determined by qPCR in MCF-7 cells left untreated or exposed to heat shock conditions for 1 hour. Data was normalized to β-actin levels. **(E)** Endogenous c-myc was detected by immunostaining in MCF-7 cells pre-treated with the indicated compounds (20µM, 1-hour) prior to a 30-minute heat shock treatment (43°C). **(F-G)** Levels of the c-myc protein were assessed by western blotting, using GAPDH as a loading control. (F) MCF-7 cells were exposed to 43°C conditions for the indicated times, or (H) heat shock treated (1 hour) cells were pre-treated (1 hour) with PI3K (Pilaralisib) or GSK3 (LY290314) inhibitors (20µM). **(H)** Schematic representation of the human rDNA cassette. Minimal E-box elements (CANNTG) in the expanded region are noted with red circles, and ChIP amplicons are noted as P1 and P2 (cyan boxes). **(I)** ChIP of c-myc from MCF-7 cells, pre-treated (1-hour) with DMSO (dark grey bars) or Pilaralisib (20µM - white bars). Data is presented as the % input obtained with the c-myc antibody in the heat shock (30 min) sample relative to the % input obtained from the untreated (DMSO) lysate. Quantification was performed by qPCR. Individual data points are included on the graphs (black or grey dots) and values on graphs represent means ± SEM (n=3). *P≤0.05 (two-tailed unpaired student’s t-test). Dashed white circles in microscopy images represent nuclei. Scale bars: 10µm.

## Discussion

In this study we used a high-throughput image-based screening approach to identify cellular factors that mediate the heat-inducible formation of functional amyloid aggregates. Our data demonstrates a role for several well-established signaling (PI3K, mTORC2, AKT, GSK3) and effector (c-myc) molecules in this process, as we generated a comprehensive cascade of cellular regulators that drive A-body biogenesis. Our analysis shows that while activation of the PI3K pathway occurred at the same temperature as A-body formation (**Figure 2A**), the temporal kinetics of AKT phosphorylation and c-myc nucleolar accumulation preceded the expression of the nucleating rIGS_16_RNA transcripts (**Figure 3A**). Considering the data demonstrating that chemical and siRNA disruption of pathway components reduces both A-body formation and rIGS_16_RNA expression (**Figure 2-4**), the most logical regulatory model for this work places the PI3K/mTORC2/AKT/GSK3/c-myc signaling proteins upstream of these ncRNA transcripts in this cellular stress response pathway.

One of the most fascinating features of this work is noting how established cell proliferation factors^52,55,56^ are regulating the formation of condensates that induce a state of cellular dormancy^28^. Activation of PI3K, mTORC2, and AKT are typically associated with increased DNA synthesis, cell cycle progression, and cell division, but under harsh environmental conditions the detention of essential cellular factors in the A-body (e.g., POLD1, cdk1, and CDC73) has been shown to arrests these same proliferative events^27,28,30,57^. How activation of the PI3K signaling axis diverges to produce dramatically dissimilar biological outcomes has yet to be established. Invariably, there are undiscovered signaling components and crosstalk molecules that drive the divergent outcomes, and further mapping of the nuances in these regulatory networks is needed in the future. At the top of the signaling cascade, it is unclear whether our screen identified the direct sensor(s) of elevated temperature. Most heat shock research focuses on the transcriptional regulator heat shock factor 1^58–60^, but there is little evidence that this molecule can serve as the receptor or adaptor molecule needed for PI3K activation. As binding to stimulated membrane receptors triggers activating conformational changes in heterodimeric PI3K complexes that expose catalytic domains^61–63^, it is conceivable that temperature-induced conformational changes could regulate PI3K proteins in a similar manner. Our recent work demonstrating that thermo-sensitive structural domains can trigger localized protein denaturation at elevated temperatures^32^, gives credence to this possibility. Thus, it would be interesting to assess whether specific members of the PI3K enzyme family have the capacity to directly sense these harsh environmental conditions.

The role of c-myc in cell dormancy and amyloid aggregation is also a striking observation. This potent proto-oncogene is frequently dysregulated in many cancer types^56,64–66^, driving classic tumorigenic phenotypes, such as proliferation, angiogenesis, and immune system suppression^67–69^. Previous work has shown that A-body formation occurs in the cores of tumors, where the hypoxic/acidotic microenvironment triggers this amyloidogenic process^28^. Without A-bodies, these tumor cores were unable to enter a dormant state, causing significant cell death within the region^28^. A role for c-myc in this process would further diversify the oncogenic functions of this protein, demonstrating that within a single tumor this transcription factor could simultaneously be promoting and impairing cell proliferation. How c-myc regulates these divergent properties has yet to be established. An association with E-box elements has been shown to control c-myc- mediated expression^70,71^, but as these enhancers are spread throughout much of the human genome there would need to be a mechanism driving the preferentially binding of c-myc to one site over another. Our data demonstrates that under heat shock conditions c-myc binding to the rIGS_16_RNA locus enhances by ∼3-4-fold (**Figure 4I**). Part of this increase could be associated with the nucleolar accumulation of c-myc protein observed during stress treatment (**Figure 4A**). A growing volume of literature suggests that this localized rise in steady state levels could be due to decreased degradation of the nucleolar population, rather than enhancing the targeting of c-myc to this subnuclear condensate^72–74^. This would be in line with our findings, as GSK3-mediated phosphorylation has been shown to increase binding of c-myc by Fbw7γ, a substrate recognition factor for the SCF ubiquitin ligase that is localized to the nucleolus^72^. Additionally, many epigenetic regulators were found to increase A-body formation (**Figure S3C**), suggesting that accessibility of the rIGS_16_RNA-encoding locus may be a factor in heat-induced c-myc binding (**Figure 4I**). Further work needs to be done to validate these proposed mechanisms, but it provides a satisfying conceptual framework for how the PI3K signaling axis can drive rIGS_16_RNA transcription and amyloidogenesis.

Overall, this study identifies a role for PI3K, mTORC2, AKT, GSK3, and c-myc in promoting the formation of functional amyloid aggregates. Our work highlights the adaptability and plasticity of this signalling cascade, as it provides an example of a cellular network whose activation can induce either proliferative or dormancy-associated phenotypes under divergent environmental conditions. It is also interesting to note that patients with Alzheimer’s disease show increased PI3K signaling in their neurons, which can positively contribute to the formation of amyloid deposits^75–78^. The parallels between PI3K-mediated activation of A-body biogenesis and pathological amyloid aggregate formation may hint at the etiological underpinnings of this neurodegenerative disease, but further study in this area is needed.

## Materials and Methods

### EXPERIMENTAL MODEL AND STUDY PARTICIPANT DETAILS

MCF-7 and U-87 MG cells were purchased from ATCC and cultured in DMEM with 10% fetal bovine serum and 1% penicillin-streptomycin, in 5% CO_2_ at 37°C. MCF-7(ATS-GFP) cells were generated by stably incorporating the previously described pF-VHL(recACM)-GFP plasmid^28^. Small molecule treatments were performed at a concentration of 20µM, for either 1-hour or 24- hours, as indicated in the figure legends. Heat shock treatments were administered by placing cells at 43°C, 5% CO_2_ for the times indicated in the figure legends. Acidosis treatments were administered in a H35 HypOxystation at 37°C in a 1% O_2_, 5% CO_2_, and N_2_-balanced environment (Don Whitley Scientific, Frederick, MD, USA) following the replacement of standard growth medium with serum-free medium at pH 6.0. Transcriptional and proteotoxic stress (TPS) treatments were induced by adding actinomycin D (ActD) (4µM) and MG132 (8µM) to the media.

### METHOD DETAILS

#### Screen Workflow

All screening was performed using the screening platform at the SFU Centre for High Throughput Chemical Biology (SFU HTCB). Glass-bottomed black walled TC-treated 384-well plates (Corning) were coated with 0.05mg/mL Poly-D-Lysine (ThermoFisher), then seeded with MCF- 7(ATS-GFP) cells using a Multidrop Combi Reagent Dispenser (2250 cells/well). The day following seeding, the SFU HTCB TargetMol L4000 library (4413 compounds) was dispensed into each plate at a final concentration of 20µM using a Tecan Evo 150 V&P Scientific with pin tools. 24-hours post-treatment, positive control wells were fixed with ice-cold methanol, then plates were heat shocked for 2-3-hours at 43°C, 5% CO_2_. The remaining wells were fixed and stained with Hoechst 33342 using a Multidrop Combi Reagent Dispenser. 6-9 frames on both DAPI and FITC channels were imaged per well (Molecular Devices: ImageXpress microscope). Images were analyzed using the multi-wavelength cell scoring module of MetaXpress software. The DAPI channel was used to detect nuclei, then A-bodies were detected on the FITC channel as any foci between 1-4µM in diameter with variable intensities (50-200 units) above background. Cells with at least 1 A-body detected were scored as positive. Wells with less than 50 total cells were removed from the analysis. The average percentage of A-body-positive cells were calculated for each well and compared to the average of all the control wells for each individual plate. Hits were identified based on individual plate controls. Wells with a score 3 standard deviations above or below the average of the negative control wells of their plate were classified as hit compounds. All wells categorized as “hits” using this threshold were verified by visual inspection. Z’ scores were calculated based on the averages and standard deviations of all the positive and negative controls for each plate.

#### Cell Fixation and Staining

Cells were fixed in either 4% formaldehyde for 20 minutes, or methanol for 5 minutes. Congo red staining was performed on formaldehyde fixed cells as previously described^28^. Immunohistochemistry was performed on methanol-fixed slides with anti-CDC73 (ThermoFisher, PA5-26189), MDM2 (Santa Cruz, sc-13161), RPA194 (Santa Cruz, sc-48385), Fibrillarin (Millipore Signam, MABE 1154; Santa Cruz, sc-25397), or c-myc (abcam, ab32072) antibodies, following established methodologies^30^.

#### RNA extraction, cDNA generation, quantitative PCR, and plasmids

Double TRIzol extractions were performed on cells treated as indicated in the figure legends, to minimize the DNA contamination within the samples. cDNA was generated from 1µg of total RNA using the qScript^TM^ Flex cDNA Synthesis Kit (Quantabio) with random primers according to the manufacturers protocol. Quantitative PCR was performed as previously described^30^. Wild- type GSK3α and GSK3β were amplified from MCF-7 cDNA by RT-PCR, and cloned into the pcDNA3.1(+). Amplification primers contained C-terminal HA-tags.

#### siRNA and plasmid transfections

30pmol of siRNA targeting AKT1 (ThermoFisher, s659), mTOR (ThermoFisher, s603), c-myc (ThermoFisher, s9129), or control (ThermoFisher, *Silencer*^TM^ Select Negative Control No.1 siRNA) was transfected into MCF7 cells using Lipofectamine^TM^ 3000 reagent (Invitrogen) 48- hours before treatment. Plasmid transfections were performed using 0.5µg of each plasmid and Lipofectamine^TM^ 3000 (Invitrogen) according to manufacturers instructions. Cells were incubated for 24-hours following plasmid transfection, prior to treatment.

#### ChIP

Chromatin immunoprecipitation was performed by adapting an existing protocol (Abcam, Cross- linking Chromatin Immunoprecipitation). MCF-7 cells treated as indicated in figure legends were incubated with 0.75% formaldehyde for 10 minutes, then quenched with 125mM glycine for 5 minutes. Following washing with PBS, cells were harvested in ChIP lysis buffer (50mM HEPES-KOH pH 7.5, 140mM NaCl, 1mM EDTA pH8, 1% Triton X-100, 0.1% Sodium Deoxycholate, 0.1% SDS, 1mM PMSF), then sonicated 9 times for 15 seconds at 40% amplitude (Branson SFX 550 Digital Sonifier). Lysates were cleared by centrifugation at 8000xg for 10 minutes, and pre-cleared with IgG (Abcam, ab37415) for 4-hours before immunoprecipitation. Anti-myc antibody (Abcam, ab32072) was incubated with lysates overnight. Pierce^TM^ Protein A/G Magnetic Beads (ThermoFisher) were added for 4-hours, prior to washing with low-salt buffer (0.1% SDS, 1% Triton X-100, 2mM EDTA, 20mM Tris-HCl pH 8.0, 150mM NaCl), high-salt buffer (0.1% SDS, 1% Triton X-100, 2mM EDTA, 20mM Tris-HCl pH 8.0, 500mM NaCl), and LiCl buffer (0.25M LiCl, 1% NP40, 1% Sodium Deoxycholate, 1mM EDTA, 10mM Tris-HCl pH 8.0). Elution buffer (1% SDS, 100mM NaHCO_3_) was added, then samples were incubated overnight in RNase A and 0.2M NaCl to reverse crosslinks. Following a 1-hour Proteinase K treatment, samples were purified using a Qiagen PCR purification kit. 4% of each elution was used per well for DNA quantification by qPCR. Primers are listed in **Table S1**, with primers for UCP2 and MXD3 promoters previously described^79^.

#### Western Blots

Western blotting was performed using standard methods, with 1:1000 dilutions of GAPDH (Santa Cruz, sc-47724), AKT (Cell Signaling Technology, 9272), pAKTS473 (Cell Signaling Technology, 9271), c-myc (Abcam, ab32072), GSK3α (Cell Signaling Technology, D80E6), GSK3β (Cell Signaling Technology, D5C5Z), pGSK3α (Cell Signaling Technology, 36E9) and pGSK3β (Cell Signaling Technology, D85E12) primary antibodies, and 1:10000 dilutions of secondary antibodies.

#### Microscopy

Quantifications of the percentages of Congo red-positive cells were based on images taken on an EVOS FL Auto 2 fluorescence microscope (ThermoFisher) with 20x objective. All other microscopy images were acquired using a Zeiss LSM880 laser scanning microscope (Carl Zeiss Microscopy, Germany) with Airyscan and ZEN 2.3 software (Carl Zeiss Microscopy, Germany). Relative A-body intensity quantifications were performed as previously described with ImageJ (National Institute of Health, USA), and quantification of the relative A-body intensity (I_r_) was using the formula I_r_ = (I_A_-I_b_)/(I_n_-I_b_), where I_r_ is the relative A-body intensity, and I_A_, I_n_, and I_b_, are the intensity of the A-bodies, nucleus, and background, respectively ^31,32^. Graphical representations depict the average relative intensity of 10 cells per sample, in 3 independent replicates.

### QUANTIFICATION AND STATISTICAL ANALYSIS

All graphs represent the mean values of at least three biological replicates, with individual data points overlaid on the bar graph. All sample sizes used in the experiments were in line with those reported in the literature for similar experiments. Error bars represent the standard error of the mean (s.e.m.), and p values were calculated using the two-tailed unpaired Student’s t-test, with the significance level of p≤0.05. Representative microscopy and western blot images were captured from experiments that were independently repeated at least three times with similar results.

### KEY RESOURCES TABLE

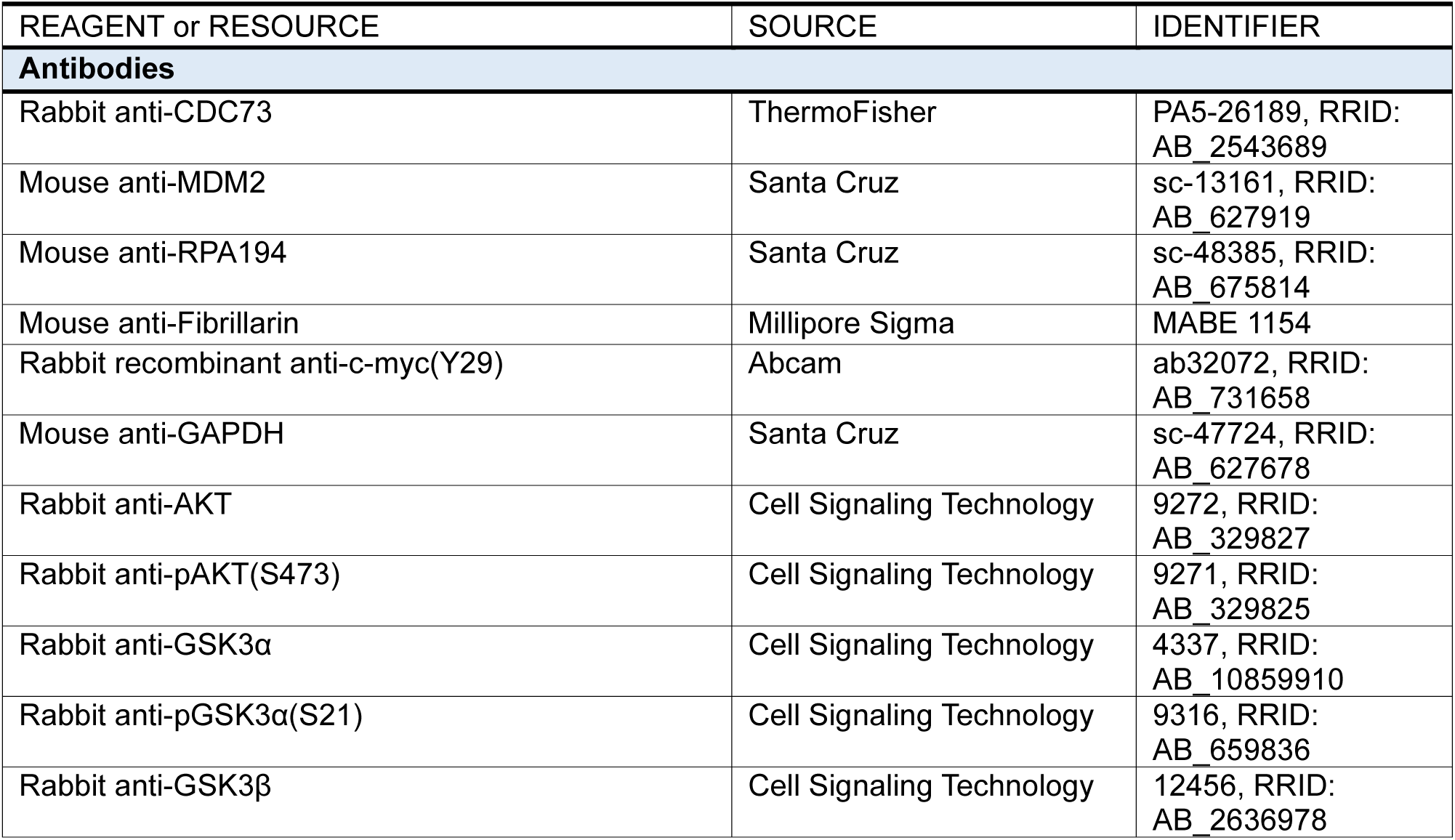
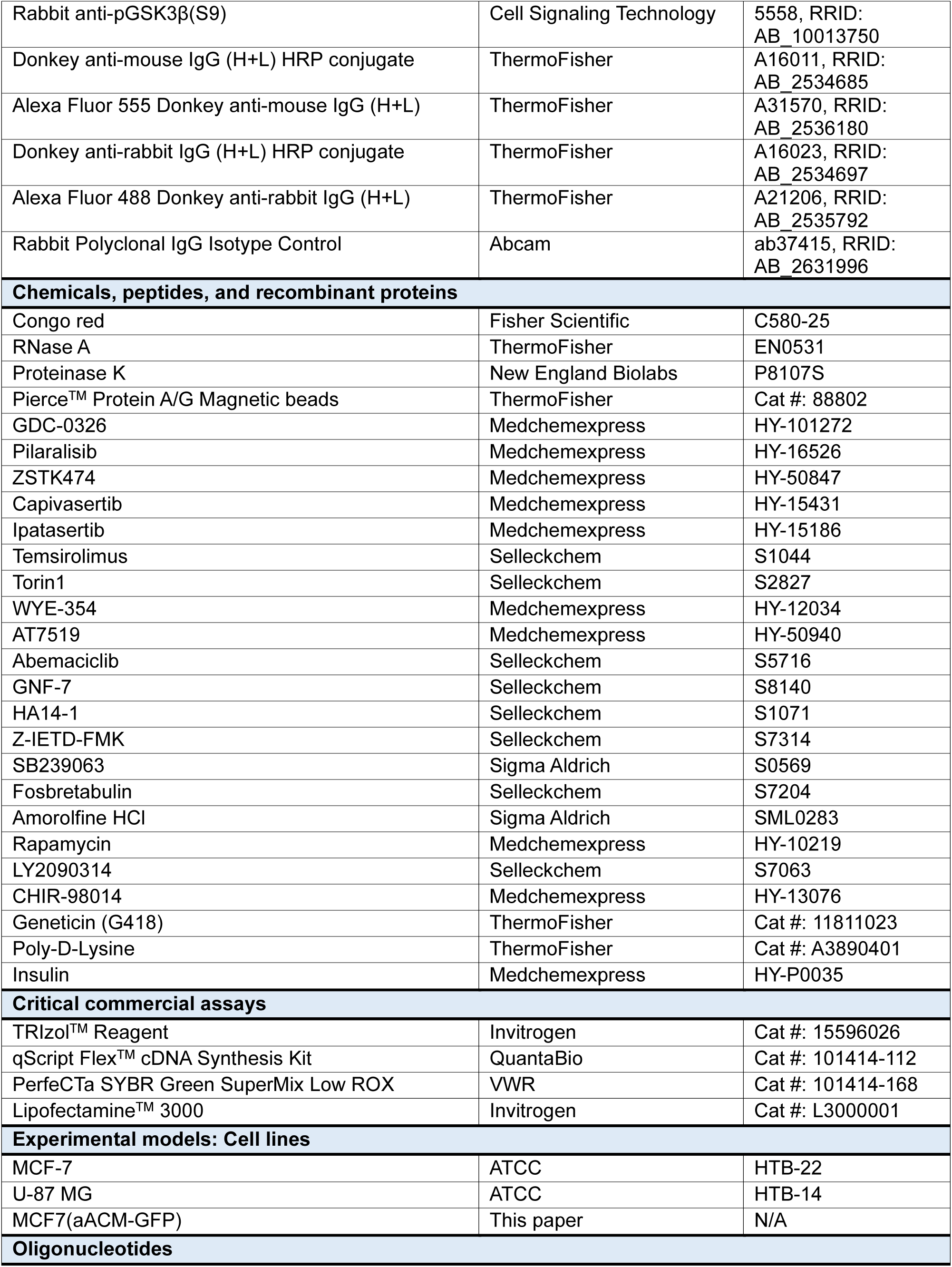
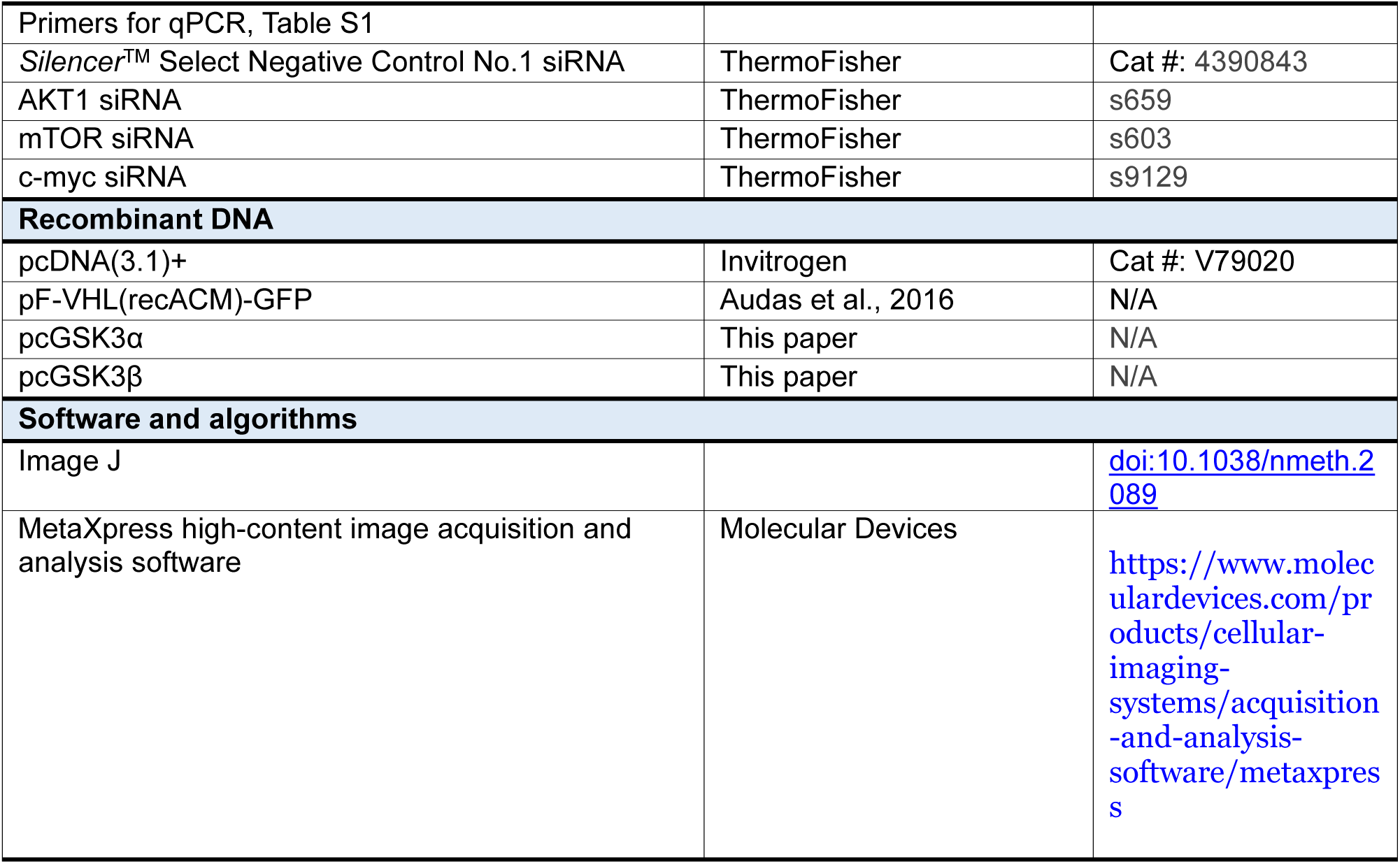

## Acknowledgements

This work was supported by the Canadian Institute of Health Research (TEA, PJT-162364) and Natural Sciences and Engineering Research Council (TEA, RGPIN-2024-04157). TEA acknowledges the kind support of the Canada Research Chairs program for a Tier II Canada Research Chair: Cellular Stress (CRC-2021-00117).

## Author Contributions

EL performed experiments, with assistance from EAM, SC, MP, and RZ. EL and TEA conceptualized the study and wrote the manuscript. All authors contributed to the review and editing. TEA supervised the project.

## Supplementary Legends

**Figure S1:**
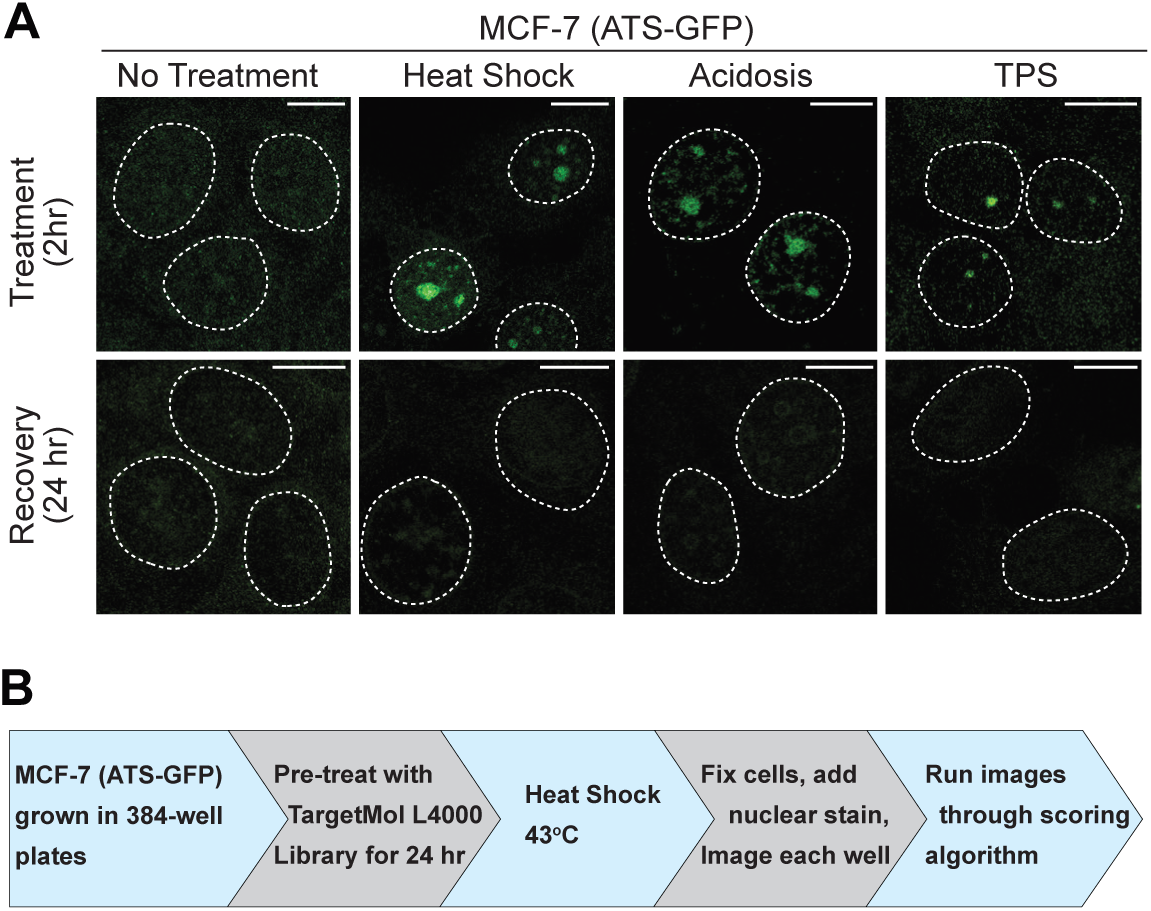
MCF-7(ATS-GFP) Cells Facilitate the Identification of A-body-Regulators. **(A)** MCF-7(aACM-GFP) cells display reversible A-body formation in all known A-body-inducing conditions. Reporter cells were treated for 2 hours with the three established A-body-inducing conditions, heat shock (43°C), acidosis (pH 6.0, 1%O_2_), and transcriptional/proteotoxic stress (TPS; 4µM actinomycin D, 8µM MG132). Following stress treatment cells were allowed to recover for 24 hours. Representative images are presented, with dashed white lines denoting nuclei. Scale bars: 10µM. **(B)** Schematic representation of image-based high-throughput screening workflow.

**Figure S2:**
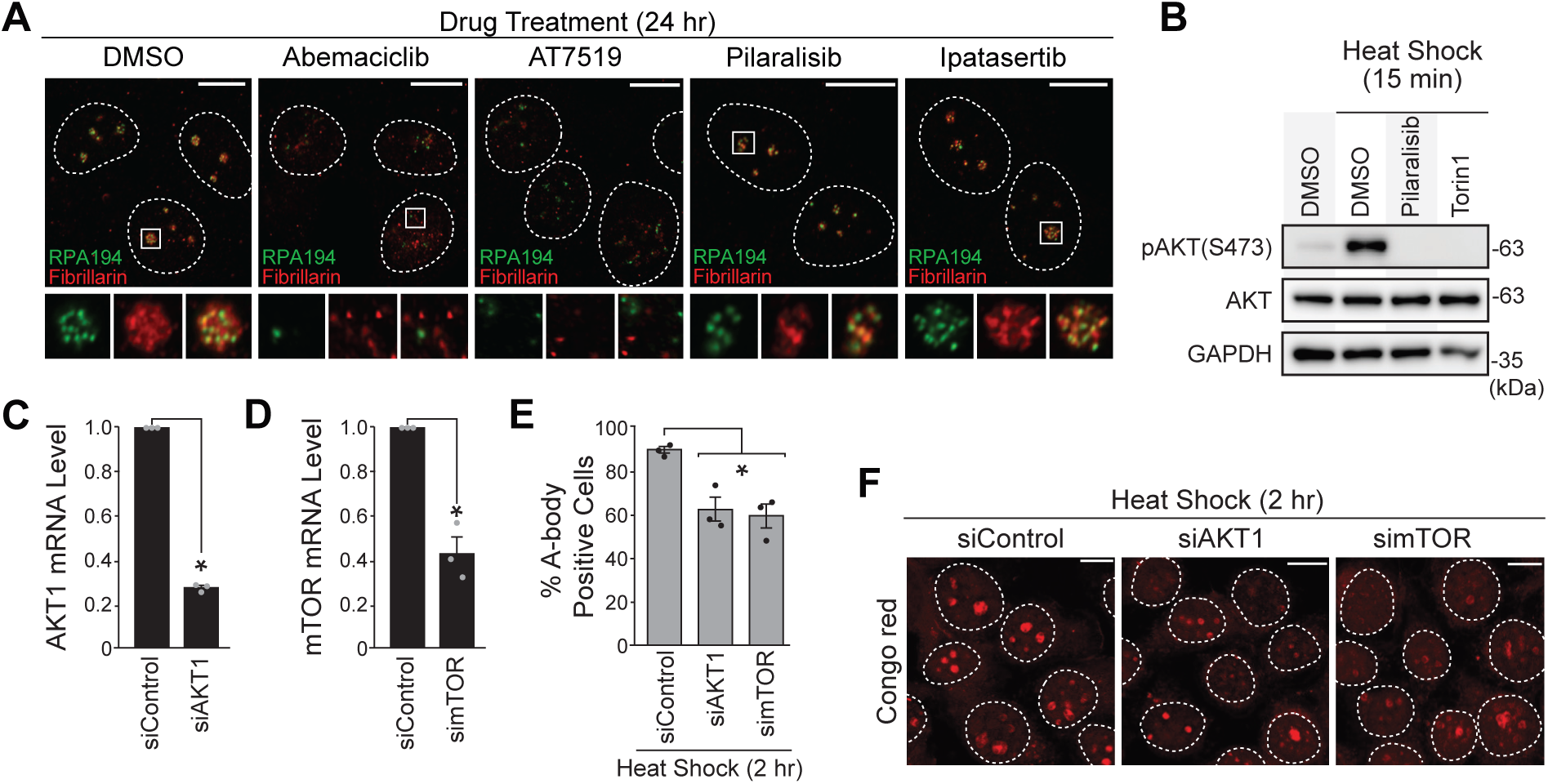
A-body Formation is Impaired by AKT1 or mTOR Depletion. **(A)** CDK inhibitors, but not PI3K/AKT inhibitors, disrupt nucleolar architecture. MCF-7 cells subjected to 24-hour pre-treatments with the indicated compounds (20µM) were immunostained to detect the endogenous nucleolar proteins fibrillarin (red), and RPA194 (green). White square insert is magnified below, with merge panel on the right. **(B)** Short pre-treatments with PI3K/mTORC1/2 inhibitors prevent AKT activation in heat shock. Total AKT, phosphorylated AKT (S473), and GAPDH were detected in MCF-7 cells by western blotting following a 1-hour pre-treatment with the indicated compounds (20µM), and a 15-minute heat shock. **(C-F)** AKT1 and mTOR depletion by siRNA reduces A-body formation in MCF-7 cells. Relative expression of AKT1 (C) and mTOR (D) mRNA in cells treated with the indicated siRNAs (30pmol, 48-hours) measured using qPCR. (E) Quantification of the percentage of A-body-positive MCF-7 cells following transfection with the indicated siRNAs and a 2-hour heat-shock. A minimum of 500 cells across 3 independent biological replicates were scored as either A-body-positive or -negative. Representative images are shown in panel (F). RNA expression qPCR data is normalized to β-actin levels. Individual data points are included on the graphs (black or grey dots) and values on graphs represent means ± SEM (n=3). *P≤0.05 (two-tailed unpaired student’s t-test). Dashed white circles in microscopy images represent nuclei. Scale bars: 10µm.

**Figure S3:**
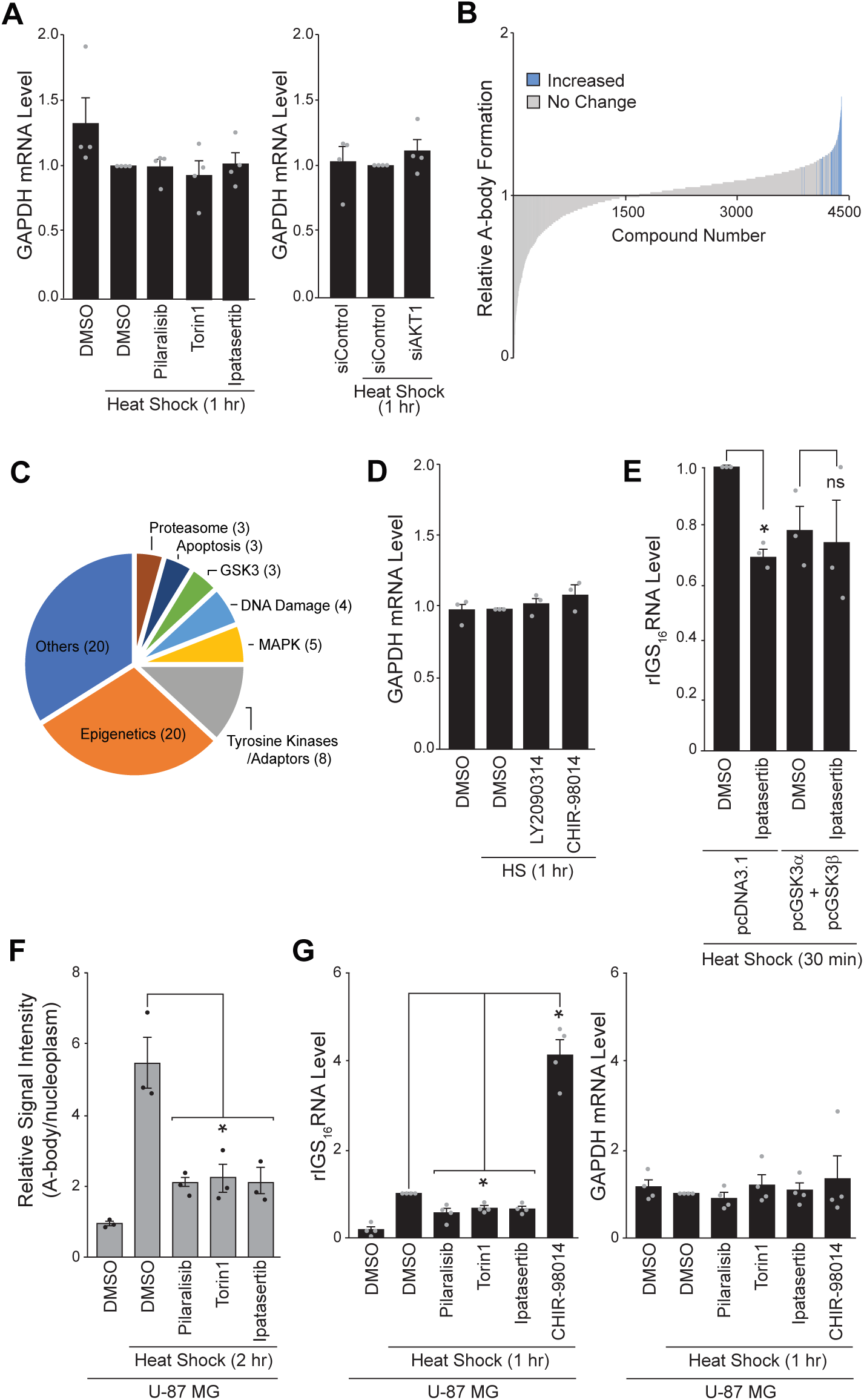
GSK3 Regulates A-body Formation. **(A)** GAPDH mRNA levels for samples from Figure 3B (left) and Figure 3C (right). **(B)** Relative A-body formation was calculated for all compounds used in our library screen that met our selection criteria. Compounds that increased A-body formation by more than 3 standard deviations from the negative control are colored blue. **(C)** Pathway analysis of screening hits that increase A-body formation. Pathways with 3 or more compounds that increasing A-body formation are listed. **(D)** GAPDH mRNA levels for samples from Figure 3I. **(E)** GSK3α/β overexpression attenuates the effects of AKT inhibition. MCF7 cells transfected with the indicated plasmids (24 hours) were treated with DMSO or Ipatasertib (20µM, 1-hour) prior to a 30-minute heat shock exposure. Relative expression of rIGS_16_RNA was measured by qPCR. **(F)** The PI3K pathway affects A-body formation in U-87 MG cells. A-body signal, relative to the nucleoplasmic background was calculated. U-87 MG cells were pre-treated with the indicated compounds (20µM, 1 hour), then exposed to heat shocked conditions (2 hours). A-bodies were detected with Congo red. **(G)** The PI3K pathway regulates the expression of the seeding rIGS_16_RNA in U-87 MG cells. qPCR analysis of rIGS_16_RNA (left) and GAPDH (right) transcript levels in heat shock treated (1 hour) cells that were pre-treated with the indicated inhibitors (20µM, 1 hour). RNA expression qPCR data is normalized to β-actin levels. Individual data points are included on the graphs (black or grey dots) and values on graphs represent means ± SEM (n=3). *P≤0.05 (two-tailed unpaired student’s t-test).

**Figure S4:**
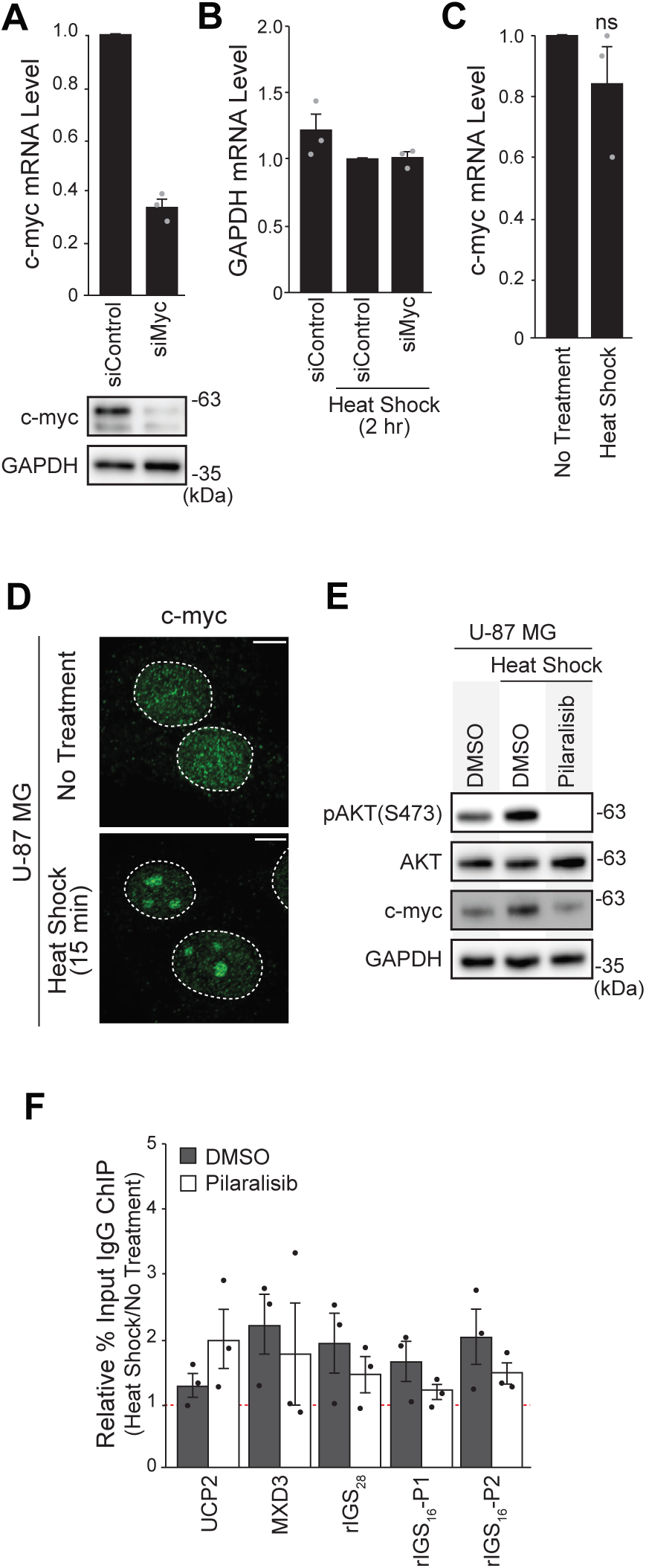
Nucleolar c-myc Promotes rIGS_16_RNA expression. **(A)** Validation of c-myc siRNA knockdown. Relative c-myc mRNA (top) and protein (bottom) levels were measured in MCF-7 cells treated with the indicated siRNA (30pmol, 48-hours) using qPCR and western blotting, respectively. **(B)** GAPDH mRNA levels for samples from Figure 4D. **(C)** Relative expression of c-myc mRNA after a 1-hour heat shock treatment. Measurements were made by qPCR. **(D)** c-myc accumulates in the nucleoli of heat shocked U-87 MG cells. Endogenous c-myc was detected by immunofluorescence analysis. **(E)** AKT phosphorylation and c-myc levels both increase in heat shock treated U 87-MG cells. Western blotting of AKT, phosphorylated AKT(S473), c-myc and GAPDH was performed in DMSO or Pilaralisib-treated (20µM, 1-hour) cells before a 1-hour heat shock (43°C). **(F)** Enrichment of IgG binding for Figure 4I. RNA expression qPCR data is normalized to β-actin levels. Individual data points are included on the graphs (black or grey dots) and values on graphs represent means ± SEM (n=3). *P≤0.05 (two-tailed unpaired student’s t-test). Dashed white circles in microscopy images represent nuclei. Scale bars: 10µm.

**Table S1:**
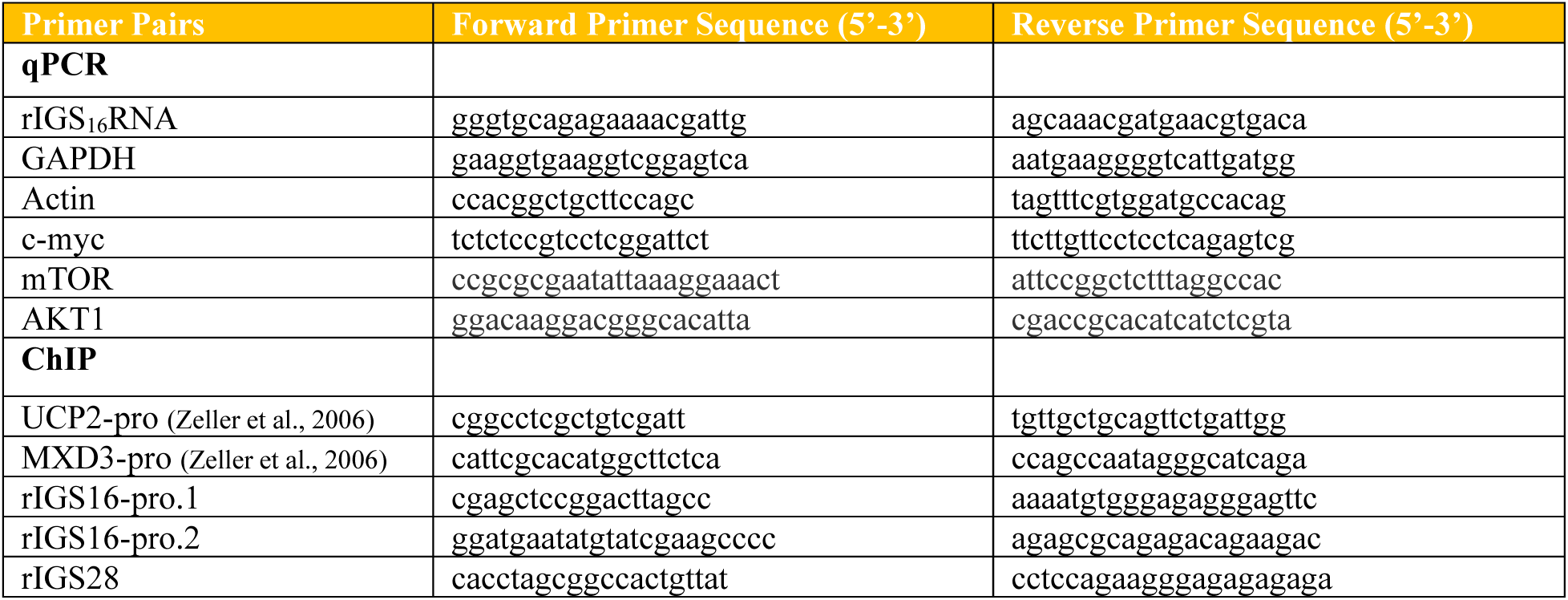
List of Primer Sequences.

